# Tick-borne flavivirus exoribonuclease-resistant RNAs contain a ‘double loop’ structure

**DOI:** 10.1101/2024.04.14.589432

**Authors:** Conner J. Langeberg, Matthew J. Szucs, Madeline E. Sherlock, Quentin Vicens, Jeffrey S. Kieft

## Abstract

*Flaviviridae* viruses are human pathogens that generate subgenomic noncoding RNAs during infection using structured exoribonuclease resistant RNAs (xrRNAs) that block progression of host cell’s exoribonucleases. The structures of several xrRNAs from mosquito-borne and insect-specific flaviviruses have been solved, revealing a conserved fold in which a ring-like motif encircles the end of the xrRNA. However, the xrRNAs found in tick-borne and no known vector flaviviruses have distinct characteristics and their 3-D fold was unsolved. To address this, we identified subgenomic flaviviral RNA formation in the encephalitis-causing tick-borne Powassan Virus. We characterized their secondary structure using chemical probing and solved the structure of one of its xrRNAs using cryo-EM. This structure reveals a novel double loop ring element leading to a model in which the ring is remodeled upon encountering the exoribonuclease. Using bioinformatic analyses we showed that this structure is representative of a broad class of xrRNAs and defined key structural and sequence determinants of function. These discoveries reveal a conserved strategy of structure-based exoribonuclease resistance achieved through a unique topology across a viral family of key importance to global health.

## INTRODUCTION

RNA viruses present a substantial global health threat, with urbanization and climate change exposing a growing portion of the population to infection^1^. Among these, the *Flaviviridae* family contains many arthropod-borne viruses such as Zika virus, Dengue virus, yellow fever virus, West Nile virus, Tick-Borne Encephalitis virus, and Powassan virus, many of which are major human health threats^2^. The genus *Flavivirus* contains positive sense single stranded RNA (+ssRNA) viruses consisting of a single genomic RNA^2^, which contains one open reading frame flanked by untranslated regions (UTR) that comprise RNA elements essential for regulating and facilitating the viral replication cycle^3–5^.

In addition to replication of the genomic RNA, infection by flaviviruses results in smaller noncoding viral RNAs called subgenomic flavivirus RNAs (sfRNAs), which result from incomplete degradation of the viral genome by host 5′ to 3′ exoribonucleases (primarily Xrn1) (Fig. 1A)^6,7^. Specifically, structured RNA elements in the genomic RNA’s 3′ UTR, called exoribonuclease resistant RNAs (xrRNAs), halt the progression of Xrn1 at discrete locations and function independently of bound protein factors; the resultant protected RNAs are sfRNAs (Fig. 1A)^5,6,8,9^. Often, viruses contain multiple xrRNAs in their 3′ UTR leading to sfRNAs of different sizes, though whether these structures are redundant or have independent functions is unknown^10^. The sfRNAs have been implicated in several cytopathic and pathological effects such as disrupting host mRNA turnover, inhibiting RNAi, altering the host interferon response, and facilitating host switching^7,9,11–16^. No analogous xrRNA structures have been identified in cellular RNAs; this and the ubiquitous presence of xrRNAs in *Flaviviridae* make them potential targets for anti-flaviviral therapeutics^17^.

**Figure 1:**
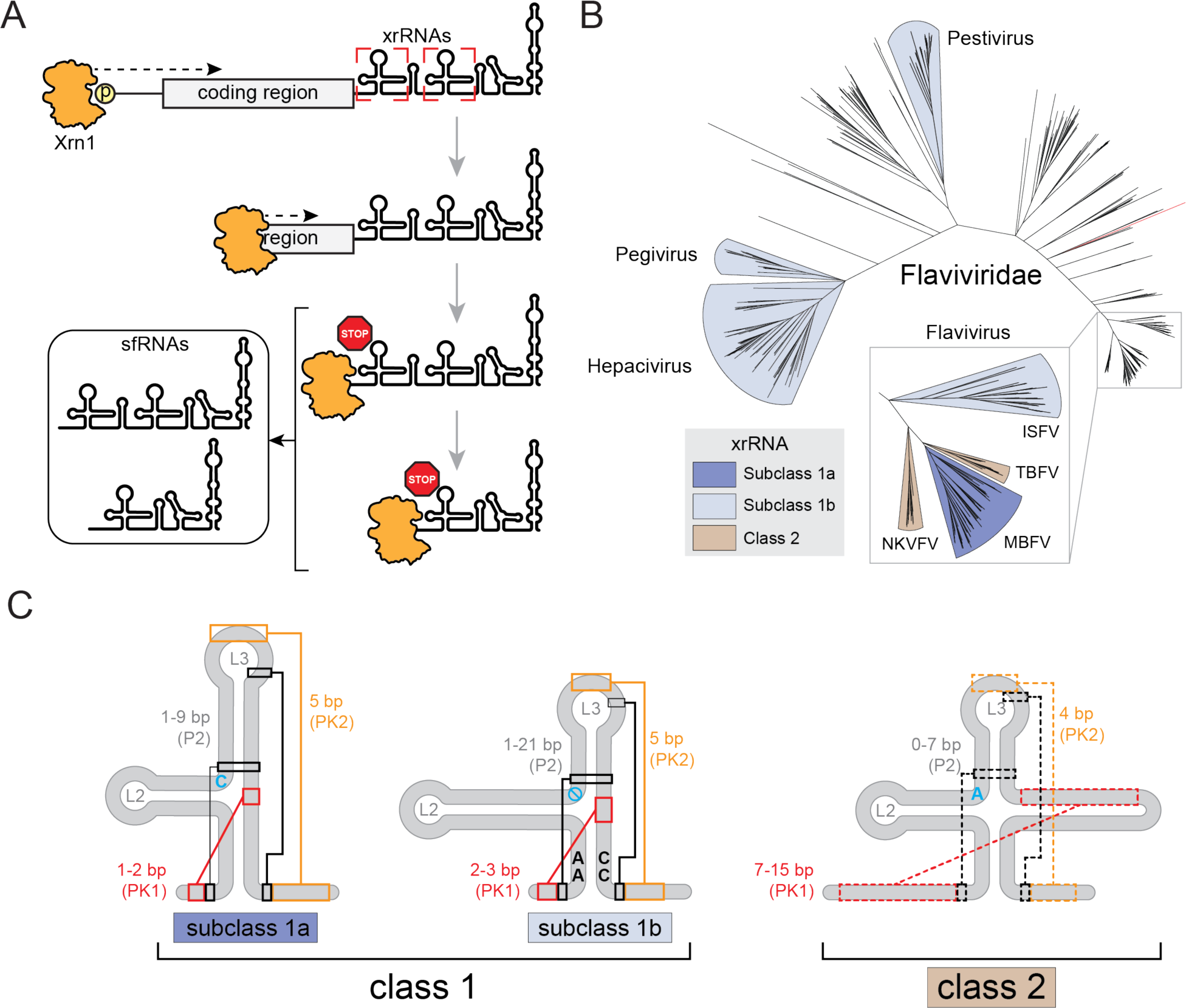
*Flaviviridae* sfRNA biogenesis and xrRNA structural determinants. **A.** A cartoon representation of sfRNA production and function through Xrn1 mediated degradation of the viral genome. **B.** Overview of the phylogenetic relationship within *Flaviviridae* based on the sequence of NS5 (RdRp). The tree files, alignments, and analysis were adapted from Bamford et al.^43^ and Mifsud et al.^44^ **C.** Structural features of *Flaviviridae* xrRNAs are depicted on the cartoon representations of the secondary structures of each class or subclass. Known long-range tertiary interaction are indicated with colored lines and boxes in the class 1. Analgous predicted interactions in the class 2 are indicated with dashed colored lines and boxes. The blue letters indicate conserved junction nucleotides, blue “no” symbol in subclass 1b denotes the absence of a nucleotide in this position.

The understanding of the structure-based mechanism of sfRNA formation comes from high-resolution crystal structures of xrRNAs from several mosquito-borne flaviviruses (MBFV) including Zika Virus (ZIKV) and Murrey Valley Encephalitis Virus (MVEV), and the distantly related Tamana Bat Virus (TABV)^5,6,8,17–20^. These xrRNAs all form a compact structure based around a three-way helical junction. The defining structural characteristic is a unique ring-like feature that encircles the 5′ end of the xrRNA. Versions of this ring-like feature have been observed in all xrRNA structures solved to date, including those from plant-infecting viruses unrelated to flaviviruses^18,19,21^. The presence of this ring-like feature combined with functional and biophysical studies show that this ring acts as a ‘molecular brace’ against the surface of the exoribonuclease, creating a mechanical block that prevents the enzyme from progressing past a defined point^5,6,22^. Tertiary interactions that stabilize flavivirus xrRNAs’ unique fold include pseudoknots, base triples, non-Watson-Crick base pairing, base stacking, and metal ion coordination. Evidence that this specific structure and stabilizing interactions are conserved among several flaviviral clades comes from comparative sequence alignments of xrRNAs found throughout *Flaviviridae*^5,6,8,9,17–19,23^.

xrRNAs are ubiquitous in *Flaviviridae* where evolution gave rise to variation in sequence and structure. Examination of covariance models of all *Flaviviridae* xrRNAs and the crystal structures mentioned above reveal differences that allowed xrRNAs to be classified: class 1 (subdivided into subclass 1a and subclass 1b) and class 2 (Fig. 1B)^17,20^. These classes are based on differences in the size, the nature of some secondary structural elements, tertiary interactions that stabilize the 3-D fold, and are supported by comparative sequence alignments. Subclass 1a comprise xrRNAs from MBFV, while subclass 1b xrRNAs are present in insect-specific flaviviruses (ISFV), TABV, and members of *Pegivirus*, *Pestivirus*, and *Hepacivirus*^17^. Although subclasses 1a and 1b have many similarities, differentiating structural features are the presence or absence of an unpaired nucleotide between P2 and P3 (Fig. 1C), and fundamental differences in the tertiary interactions that ‘close’ the ring feature^8^.

The class 2 xrRNAs lack structural characterization compared to class 1 and have thus far only been found in tick-borne flaviviruses (TBFV) and no known vector flaviviruses (NKVFV) (Fig. 1B)^20,23^. The secondary structures of the class 2 xrRNAs have been depicted several ways, but recent analysis^20^ led to versions showing features reminiscent of the class 1 with some key differences (Fig. 1C). Specifically, within the helical junction, the unpaired junction nucleotide is conserved as an A in all known sequences of class 2, while it is always a C in the subclass 1a and is always absent in the subclass 1b. Additionally, an important pseudoknot (PK1) in class 2 appears to be significantly longer (6-10 base pairs) than in class 1 (1-3 base pairs)^20,24^ (Fig. 1C).

Based on its similarities to the class 1, it is hypothesized that the class 2 xrRNAs also contain a ring-like feature important for blocking Xrn1, but there is no 3-D structural information to verify this. Likewise, it is not clear how a ring-like structure could be accommodated given the larger proposed pseudoknot structure. Also, the location where the exoribonuclease halts on class 2 xrRNAs is within a base-paired element rather than in unpaired RNA upstream of the xrRNA ring structure^23^. If class 2 contain a ring-like feature, it must have substantial deviations from the class 1, given the differences in the proposed secondary structures. In addition, bioinformatic searches for new xrRNAs would be enhanced by structural insight, as current knowledge of class 2 comes mostly from biochemical studies^20,23^.

Although we have successfully crystallized several class 1 xrRNAs, attempts to obtain diffraction-quality crystals of representative class 2 xrRNAs were unsuccessful. However, single particle cryo-electron microscopy (cryo-EM) has emerged as an invaluable method for determining the structure of biomolecules previously inaccessible by traditional structural approaches. Cryo-EM was recently applied to several RNAs to determine tertiary structure and dynamics, and we recently presented a scaffold-based approach to facilitate the structural determination of diverse RNAs by cryo-EM^25–31^. We therefore chose to apply this scaffold-based approach to the structure of a class 2 xrRNA from Powassan virus (POWV). Powassan is an tick-borne human pathogen found in both North America and north-east Asia^32^, but the presence of xrRNAs in the POWV genomic RNA and its ability to form infection-critical sfRNAs had not been verified, and the structures of any putative class 2 xrRNAs were unknown.

Here we report the results of combined virological, biochemical, bioinformatic, and structural methods applied to the xrRNAs from POWV. We verified the presence of two xrRNAs in POWV genomic RNA and show they result in sfRNA formation. We used cryo-EM to determine the structure of a POWV class 2 xrRNA, revealing the presence of a ring-like feature that is more extensive than any previously observed, forming a ‘double loop’ around the 5′ end of the xrRNA. This ring may interact with Xrn1 in unique ways compared to the ring in class 1. Structure-informed bioinformatic analysis then helped us catalog a more complete set of class 2 xrRNAs and define key features of the class. Overall, our findings complete our knowledge of the diversity of 3-D structures in *Flaviviridae* xrRNAs, and further highlight the essential role of conserved but variable ring-like folds stabilized by diverse class-specific tertiary interactions.

## RESULTS

### Powassan virus produces sfRNAs during infection

We chose to study the putative class 2 xrRNAs from the Spooner isolate from the Deer tick lineage of POWV^32^, as it is an emerging human pathogen for which no targeted therapies exist, and the sequence of its 3′ UTR suggested the presence of two class 2 xrRNAs^20,23^. To verify that POWV infection results in the production of sfRNAs, we infected Vero cells and assessed sfRNA generation by northern blotting at various time points and multiplicities of infection (MOI) (Fig. 2A). Blots revealed two discrete bands of sizes consistent with sfRNA1 and sfRNA2 species starting at the 5′ end of the first and the second putative xrRNA structures, respectively. To verify that these sfRNAs were due to authentic xrRNAs, we used an established *in vitro* Xrn1 resistance assay. Briefly, *in vitro* transcribed RNA comprising the entire POWV 3′ UTR was challenged with recombinant Xrn1 then resolved by denaturing polyacrylamide gel electrophoresis (Fig. 2B). We observed two bands of the expected sizes that would result if the enzyme began degrading the input RNA, but then halted at one of the two putative xrRNAs, matching the pattern from infection. These results not only confirm the production of sfRNAs by the xrRNAs in POWV during infection, but also demonstrate these structures function without protein cofactors, a hallmark of true xrRNAs. This observation also establishes these xrRNAs as a model for class 2 xrRNAs.

**Figure 2:**
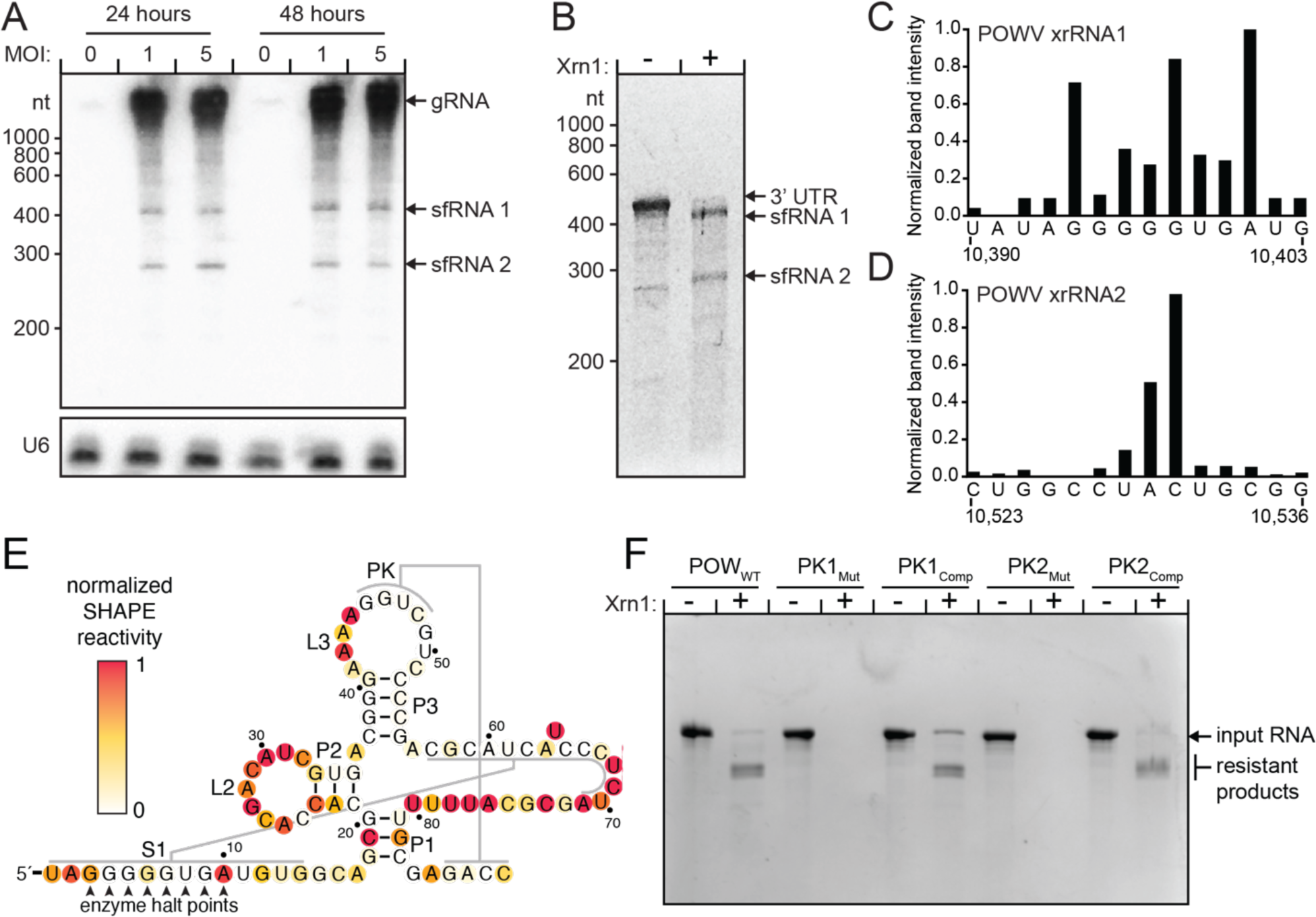
Identification characterization of POWV xrRNAs. **A.** Northern blot of POWV infected Vero cells at the indicated MOI and time post infection. Probes were specific for the 3′ UTR of POWV or the U6 RNA loading control. Infection in northern blots were confirmed in three independent experiments, one is shown. **B**. In vitro degradation by Xrn1 of in vitro transcribed RNA of the POWV Spooner isolate 3′ UTR. The partially degraded but stable RNAs appear as bands in the lane with Xrn1, demonstrating the presence of xrRNAs. **C and D**. Stop site analysis of POWV xrRNA1 and xrRNA2. RNA treated with Xrn1 was analyzed using reverse transcription primer extension. The intensity of the resultant bands is graphed as a function of nucleotide position. **E.** Secondary structure diagram of the POWV xrRNA1 in the context of the full 3′ UTR with the results of 1M7 chemical probing. **F**. *In vitro* RNA resistance assay demonstrating the functional importance of both PK1 and PK2. Diagram of the mutations is found in Supplementary Figure 3.

### The exonuclease halt site on POWV class 2 xrRNAs differs from the class 1

In class 1 xrRNAs, the Xrn1 halt site resides a few nucleotides upstream of the P1 helix and pseudoknot 1 (PK1), in unpaired 5′ nucleotides that emerge from the center of the ring-like structure^9^. In class 1, the 1-3 base pairs that comprise PK1 lie within the ring and are essentially inaccessible to the approaching enzyme. This configuration supports the idea that the ring braces against the surface of the enzyme and prevents it from unwinding base pairs that are within or behind the ring. However, in class 2, the proposed PK1 is much longer (Fig. 1C). In previous studies, the halt point on other class 2 xrRNAs was mapped to a base-paired region that would lie within this longer pseudoknot (termed P1.1/P1.2 in some previous depictions)^23^, in contrast to class 1.

To map the halt site on POWV xrRNAs, we treated *in vitro* transcribed POWV 3′UTR RNA with Xrn1, recovered the xrRNA-protected products, and mapped the resulting 5′ ends using reverse transcription (Fig. 2C, D). In contrast to the class 1 xrRNAs – where the halt site exists primarily at a single position four nucleotides upstream of PK1 – in POWV the halt sites are located within PK1 for both the first and second POWV xrRNA (Fig. 2E). This indicates that the enzyme must unwind parts of PK1 as it approaches the xrRNA, but then halt. Furthermore, we observed a ‘stuttering’ effect; that is, there appear to be several stop sites within the xrRNA rather than a well-defined single location (Fig. 2C-E). Specifically, we identified three halt sites within PK1 in the first POWV xrRNA and multiple halt sites the second POWV xrRNA, although to a lesser extent. This suggested a different configuration or role for the extended PK1 and any ring-like structure in class 2 versus class 1, or perhaps even a different mechanism of halting Xrn1.

### The POWV xrRNAs adopt a conserved secondary structure

Existing models make robust predictions regarding the secondary structures of the POWV xrRNAs, but the unusual halt sites suggested there may be features not obvious in those analyses. Also, the fact that the enzyme must partially unwind the xrRNA to reach all potential halt points raised the question of whether the xrRNA secondary structure could change in response to partial degradation. We therefore experimentally queried the secondary structure of the POWV class 2 xrRNA in different contexts using chemical probing, specifically by DMS-MaP^33^ (Dimethyl sulfate (DMS; modifies Watson crick face of A and C)^34^) and SHAPE-MaP (selective 2′-hydroxyl acylation analyzed by primer extension (1-Methyl-7-Nitroisatoic Anhydride (1m7; modifies 2’OH in flexible backbone regions)^35^ mutational profiling). To recapitulate the viral genomic context of the POWV xrRNA, we probed constructs consisting of the (1) full 3′ UTR, (2) sfRNA1, and (2) sfRNA2; the latter two being sfRNAs produced by *in vitro* treatment with Xrn1 followed by probing.

Overall, the measured reactivity profiles of the xrRNAs from the POWV 3′ UTR, sfRNA1, and sfRNA2 were all similar, indicating the secondary structure does not change in these different contexts (Fig. 2E, S1, S2). Nucleotide reactivities from both 1m7 and DMS were in agreement with features present in the proposed class 2 xrRNA secondary structure model^20^. The three-way helical junction is present in both xrRNAs, demonstrated by the overall high reactivities in the L2 and L3 loops that are not involved with tertiary interactions. The P1, P2, and P3 helices generally show low reactivity except for some of the nucleotides at the terminal end of the helix, this is expected from ‘breathing’ in terminal pairs. The large unpaired region after the proposed PK1 is accessible for modification in all constructs supporting the same secondary structure in all contexts.

Importantly, the presence of both pseudoknots was supported by both 1m7 and DMS probing. To further test their presence and importance, we made substitution mutations that disrupted either PK1 or PK2 and tested them for Xrn1 resistance in vitro (Fig. 2F, S3). Both lost the ability to block the exoribonuclease, but resistance was rescued with compensatory mutations (Fig. S3). This matches results performed on xrRNAs from other viruses^23,24^. However, elevated reactivities within the 5′ portion of PK1 suggest that while the pseudoknot forms, there are conformationally dynamics present at the 5′ end. We note that these more dynamic base pairs correspond to the location just upstream of the mapped stop sites in the xrRNAs (Fig. 2E).

### Cryo-EM yields an intermediate resolution map and model of POWV xrRNA structure

To better interpret the functional and biochemical data and understand the structural basis of exonuclease resistance in class 2 xrRNAs, we solved the structure of a POWV class 2 xrRNA using cryo-EM with our previously described scaffolding approach^31^. Briefly, the xrRNA was appended to a modified version of the *Tetrahymena* ribozyme, creating a large and structurally recognizable chimeric RNA amenable to cryo-EM. The connection point was through the xrRNA’s L2 loop, a region which is not conserved in sequence, does not make any known long-range interactions, and is known to be dispensable in other xrRNAs^5^ (Fig. S4A). The xrRNA’s L2 loop was connected to the ribozyme’s P6b element; this has been shown to not disrupt the folding of the scaffold RNA or the appended domain^31^. To ensure appending the xrRNA to the scaffold did not perturb xrRNA fold or function, we performed Xrn1 resistance assays on this scaffolded version of the POWV xrRNA. It maintained resistance to Xrn1 *in vitro*, indicating adopted the correct functional fold when appended to the large scaffold tetrahymena ribozyme RNA (Fig. S4B).

We used this RNA to prepare grids and collect cryo-EM data that was analyzed using a workflow from our previous work^31^. Following multiple rounds of 2D classification, classes were present in which the appended xrRNA domain is clearly observed (Fig. S5). The resulting 493,118 particles were then used for initial map generation. The maps had clear features corresponding to the xrRNA, revealing its global architecture. However, 3D variability analysis revealed a degree of structural heterogeneity among this set of particles. To recover the most complete map possible, we used 3-D classification to partition these states (Fig. S5). The resulting maps yielded several volumes corresponding to what appeared to be various levels of ‘melting’ of the distal regions of the structure. This supports the idea that these regions are conformationally dynamic, consistent with the chemical probing data.

Moving forward with the most well resolved and complete map of the xrRNA calculated from 55,485 particles, non-uniform refinement yielded a map with a global resolution of 4.1 Å in the scaffold domain with a local resolution of the xrRNA of 5.1 Å (Fig. 3A, S5). Within the xrRNA portion of the map, clear structural features are observable, with both major and minor grooves apparent and phosphate bumps clear along the backbone for most of the map (Fig. 3B). Using this map and information from the chemical probing and covariation analysis, we built a model of the xrRNA (Fig. 3C,D). While the local resolution of the xrRNA portion of the maps did not achieve ‘atomic level’ high resolution, all regions were of sufficient resolution to reliably place the resolved nucleotides.

**Figure 3:**
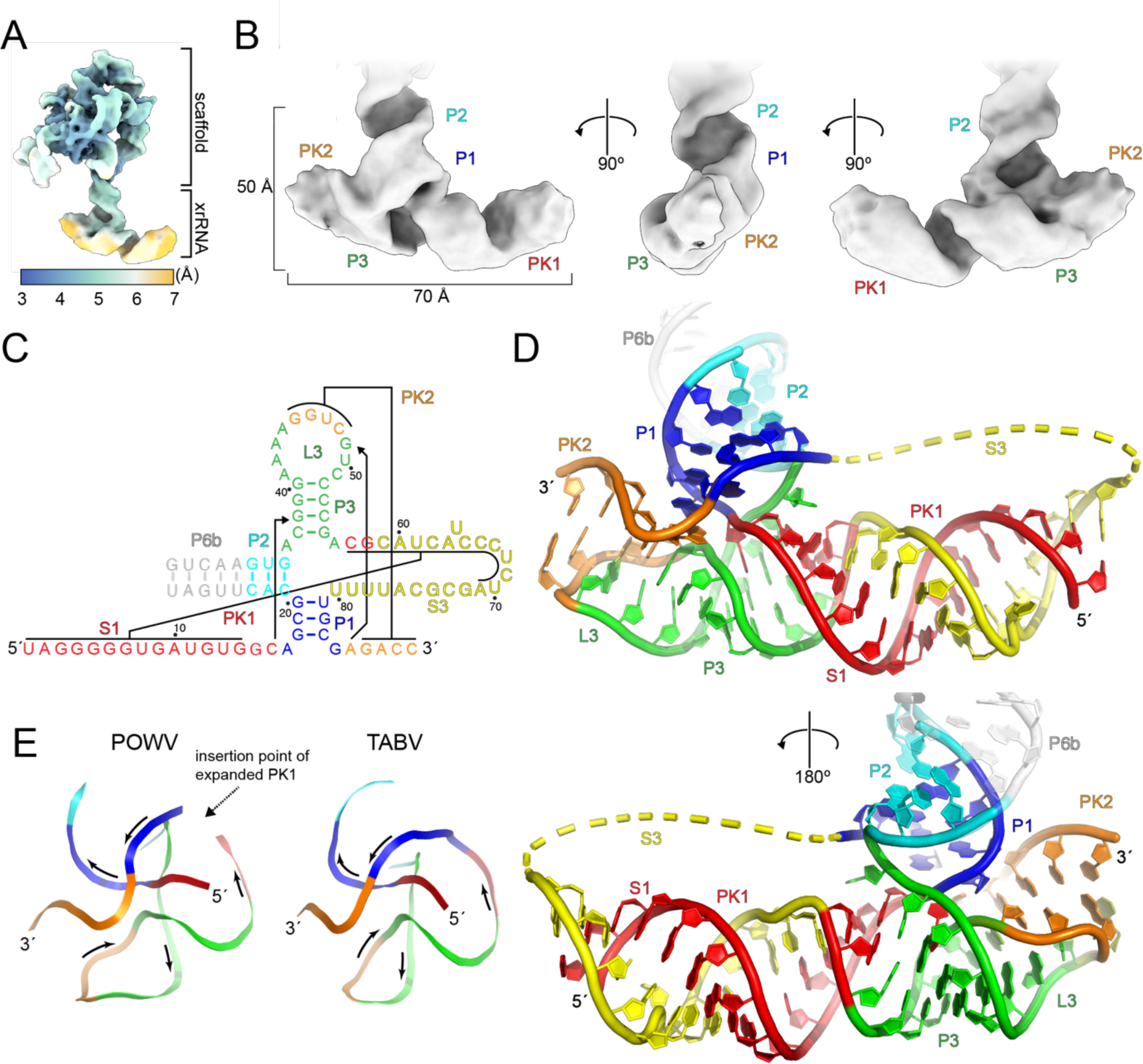
Cryo-EM structure of the POWV class 2 xrRNA. **A.** Local resolution map of the scaffolded POWV xrRNA. Resolution is colored as denoted in the inset key **B**. Cryo-EM map of the POWV xrRNA, the location of secondary structure features are indicated. **C**. Secondary structure diagram of the POWV xrRNA, colored to match panels D and E. **D**. Atomic model of the POWV xrRNA colored to match panel C. The yellow dashed backbone represents 9 nucleotides that were not visible in the map and hence not built. Structural features are labeled. **E.** Comparison of POWV and TABV ring topologies colored corresponding to panel C.

### Structure of a POWV xrRNA has features reminiscent of other xrRNAs

The POWV xrRNA structure has several features that match the class 1 xrRNAs, indicative of function dependent on a specific shared 3-D structure (Fig. 3C,D). First, the expected paired helical elements P1, P2, and P3 are present, as is pseudoknot 2 (PK2) formed through pairing between sequence at the 3′ end with part of L3. These stems adopt an overall configuration similar to class 1 xrRNAs in which P1 stacks with P2, while P3 stacks on PK2^5,6,8^. Interestingly, the overall angle between the helices and the trace of the backbone through the core of the POWV xrRNA structure are much more like the subclass 1b xrRNA from TABV than the subclass 1a from ZIKV (Fig. 3E). As with the class 1 xrRNAs, the POWV class 2 core structure gives rise to a ring-like motif, although it differs substantially from previous structures (see Discussion).

As with class 1 xrRNAs, the POWV xrRNA contains a base triple between a C nucleotide located upstream of the P1 helix and a G-C pair in stem L3 (C^+^16•G38-C54). The same triple is found in other class 1 xrRNAs, where it can be C^+^•G-C or U•A-U. In all these xrRNAs, this triple appears to be a key interaction between the 5′ end of the xrRNA and the three-way junction, positioning the 5′ end to pass through the center of the ring.

The POWV xrRNA structure also contains a single unpaired nucleotide, A36, between P2 and P3 that is involved in defining the ring-like motif through base stacking. This matches a highly conserved C nucleotide at this position in the subclass 1a xrRNA where it is a defining characteristic. While both the class 2 and subclass 1a contain a nucleotide in this position, it is distinctively absent in the subclass 1b^17,20^. Xrn1 resistance experiments testing substitutions, deletions, and insertions at this position show that each class of xrRNA is highly sensitive to the identity and presence or absence of the correct nucleotide in this position^5,6,8,9,17^. Analysis of the structures of different xrRNAs suggests this strict requirement may correlate to the size or nature of the ring-like structure and specific stabilizing interactions in different xrRNA classes^17,20^.

### Bioinformatic analyses reveals conserved determinants of exonuclease resistance

Using the 3-D structure of a POWV class 2 xrRNA as a guide, we undertook a bioinformatic analysis to (1) identify additional examples of class 2 xrRNAs, (2) develop a more robust understanding of the class 2 sequence and structural constraints, and (3) determine if the structure of the POWV xrRNA is representative of class 2 xrRNAs in general. This type of approach has proven successful in identifying novel variants of structured RNAs as well as supporting proposed folds through statistically significant covariation^36–40^. We used a comparative sequence alignment from 28 previously identified class 2 xrRNA sequences^20,23^ with the program Infernal^41^ to query a database containing all deposited viral genome sequences (retrieved 01/06/2023). This search identified 38 putative class 2 xrRNAs, 35 from TBFVs and 3 from NKVFVs (Fig. S6A). Of the 13 recognized TBFV species, 10 were identified in this search, including both lineages of POWV as well as multiple subtypes of tick-borne encephalitis virus.

We calculated a consensus sequence and secondary structure from the identified sequences using CaCoFold visualized with R2R^37,40,42^. The resulting covariation patterns supported the secondary structure elements observed in the cryo-EM structure of the POWV xrRNA, recovering all stems and pseudoknots present in this class (Fig. 4A), and showed that the helical organization and base pairing schemes present in the POWV xrRNA are general to all class 2 xrRNAs. These results largely agree with previous work bioinformatically characterizing class 2 xrRNAs^23,24^. However, with the cryo-EM structure, we were able to provide evidence for several additional structural features critical for function. Most striking is the extended PK1 and double loop. Several additional covariations-supported base pairs were identified, representing a more extensive base pairing context between the 5′ end of the structure and the seemingly unpaired sequence between P3 and P1. Indeed, this length ranges from 7 to 15 base pairs. In order to accommodate this extended interaction, the linker region between PK1 and P1 (S3) is sufficiently long in all cases to allow for the full formation of PK1 (Fig. S6B, C). Additionally, the highly conserved C^+^•G-Y base triple is verified as universal in this class. Based on the alignment, the P1 helix consists of three Watson-Crick base pairs and a terminal non-Watson-Crick pair consisting in most cases of an A•G pair. In some cases, this is a replaced by a CC dinucleotide, possibly yielding a similar pairing scheme as in subclass 1b xrRNAs^8,17,20^. If present, P2 ranges between 1 and 7 base pairs with an average of 5. The four-base pair P3 helix length is conserved with ∼92% of the aligned sequences containing a CGGG-CCCG pattern and the remaining 8% of sequences containing UGGG-CCCA or UGGG-CCUA (Fig. S6A).

**Figure 4:**
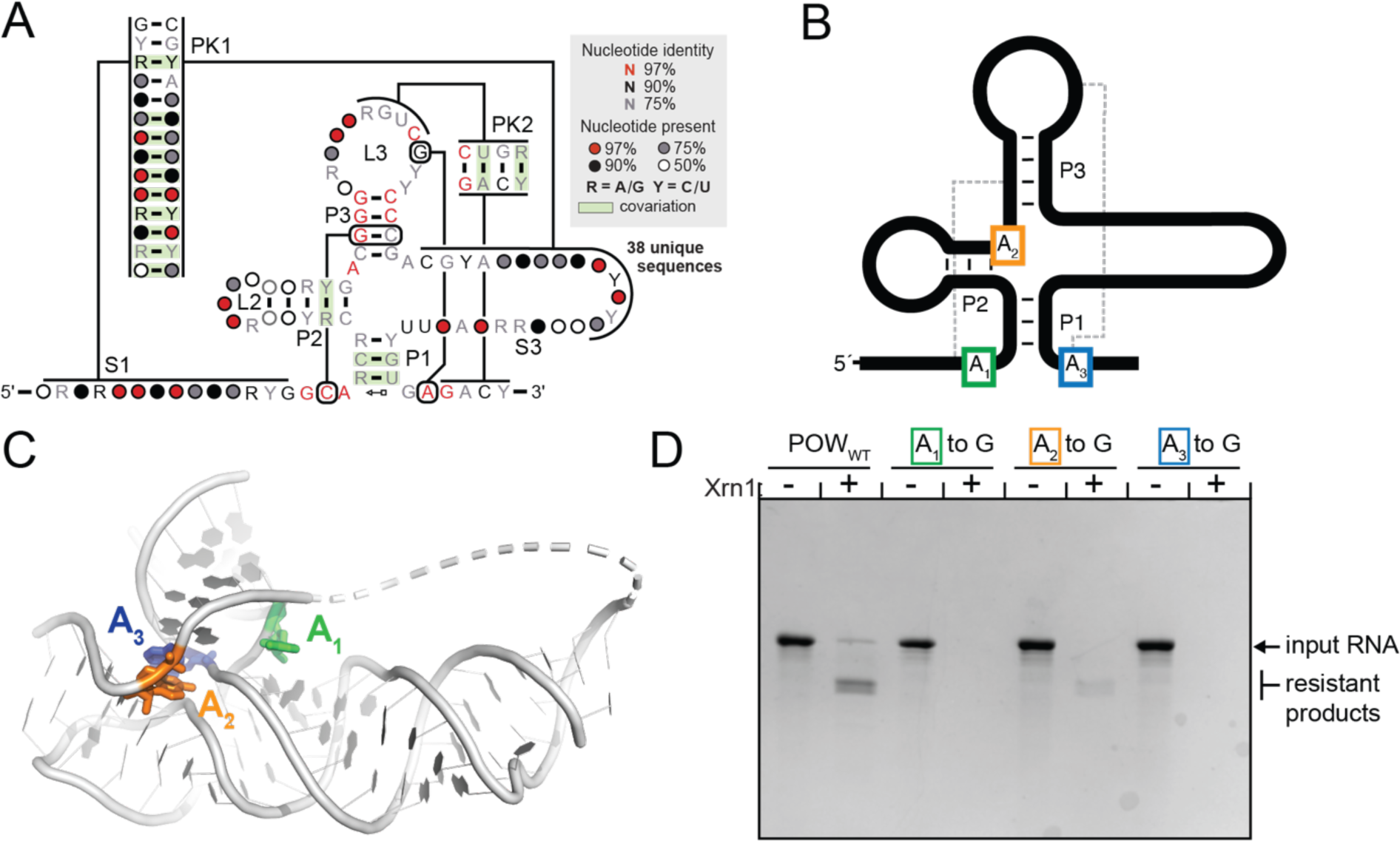
Sequence and structure conservation of class 2 xrRNAs from flavivirus. **A.** Consensus secondary structure model of class 2 xrRNAs calculated by R-Scape^37,40^. **B**. Diagram denoting the location of three conserved A bases that were mutated to test their functional significance. **C**. Structure of the POWV xrRNA with the three conserved A bases indicated in panels B and C. **D.** In vitro RNA resistance assay demonstrating the functional importance of the three conserved A nucleotides in class 2 xrRNAs.

Several perfectly conserved A bases are present in the structure, appearing to make key long-range interactions to facilitate resistance (Fig. 4A-C). As noted previously, in class 2 xrRNAs, a perfectly conserved A base is present between P2 and P3. Two others are located adjacent to P1, involved in non-Watson-Crick interactions that appear to close the ring (Fig. 4B). Analogous bases are present in other classes of xrRNA that are essential in forming the resistant motif^5,6,8^. To test the importance of these interactions in the POWV class 2 xrRNA, we prepared RNA in which each A was individually mutated to a G and challenged these RNAs with Xrn1 *in vitro*. Point mutations of any one of these bases resulted in the loss of resistance, supporting their involvement in forming the proper fold (Fig. 4D). That all three are universally conserved in class 2 xrRNAs suggests they are critical features of this class. Overall, class 2 xrRNAs appear to be well conserved structurally and our consensus model suggests the same structure-based mechanism of resistance exists across the entire class.

### The POWV class 2 xrRNA contains a novel ‘double loop’ structure

While many features of the POWV class 2 xrRNA match class 1, a dramatic difference is in the size of PK1 and the resultant expansion of the ring-like feature into a form that we refer to as a ‘double loop’ (Fig. 5A). In class 1 xrRNAs, the 2-3 base pair long PK1 forms stacking interactions with P2 and positions the 5′ end of the xrRNA to pass through the center of the fold^5,6,8^. The 13 base pair long PK1 in the POWV xrRNA also positions RNA to pass through the middle of the fold, but forms a stem that extends away from the core of the structure (Fig. 2D). In fact, the expansion of PK1 makes it the longest base paired element in the structure and the junction within class 2 flavivirus xrRNAs is technically a four-way junction between P1, P2, P3, and PK1. The 3′ end of PK1 is linked to the 5′ end of P3 by a stretch of nine unpaired nucleotides (S3) that are not visible in our cryo-EM maps and hence could not be modeled. This is likely due to a high degree of conformational flexibility in these unpaired nucleotides, consistent with the chemical probing (Fig. 2E).

**Figure 5:**
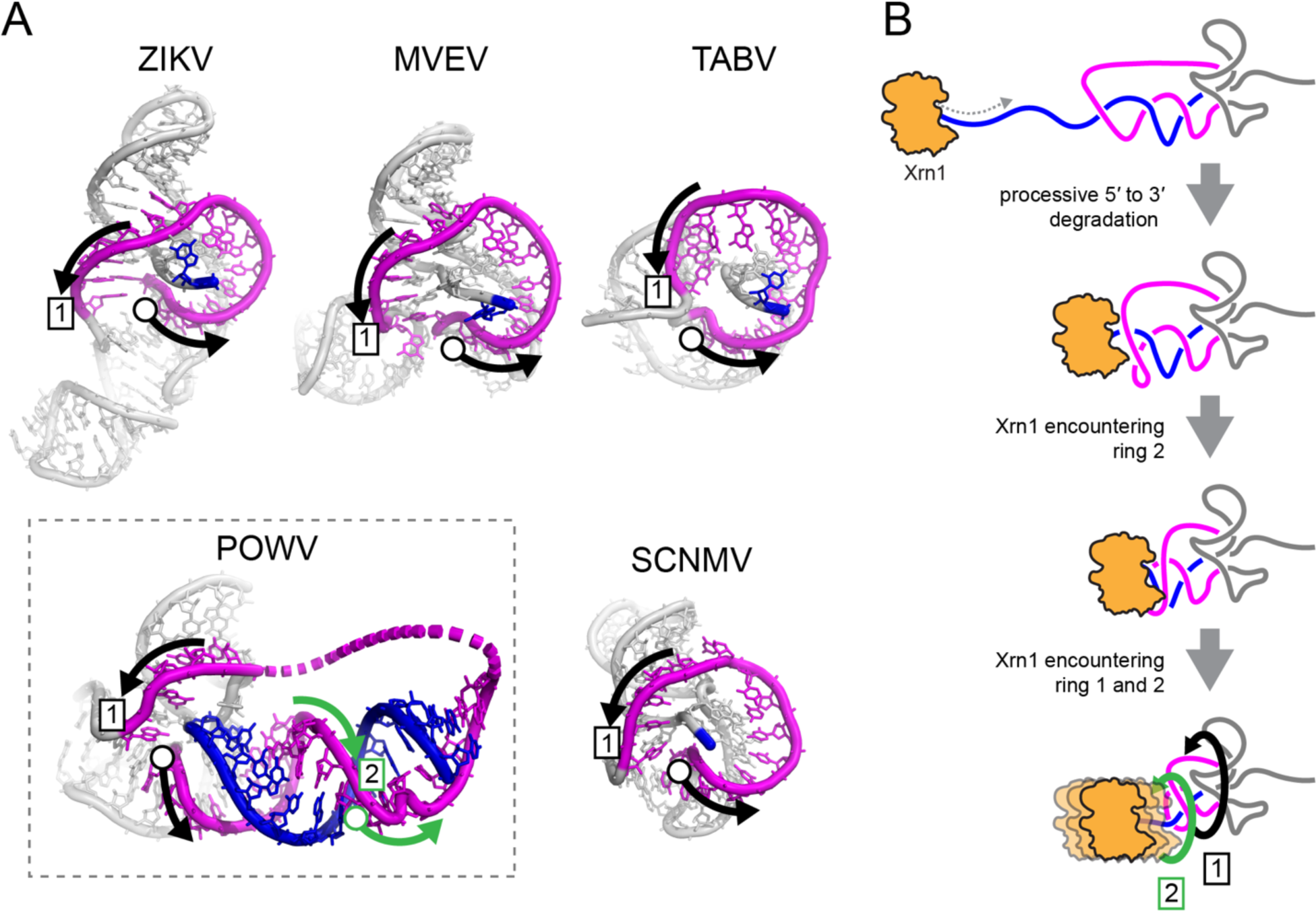
A conserved ring-like fold present across all xrRNA structures. **A.** The ring-like structure (magenta) present encircling the 5′ end (blue) of the xrRNA from ZIKV (PDB:5TPY)^5^, Murray Valley Encephalitis virus (MVE) (PDB:4PQV)^6^, TABV (PDB:7k16)^8^, POWV (this study), and Sweet Clover Necrotic Mosaic Virus (SCNMV) (PDB:6D3P)^19^. The open circle denotes the start of the ring structure, the boxed “1” indicates the completion of one loop. In POWV, the RNA loops around the 5′ end a second time, indicated with green arrows and a boxed “2.” **B.** Proposed mechanistic model of Xrn1 encountering a class 2 xrRNA based on the three-dimensional cryo-EM structure and biochemical experiments. Depicted is the ‘double loop’ consisting of the primary ring-like motif found in all solved xrRNAs to date (1) and the second ring resulting from the extended PK1 interaction in class 2 xrRNAs (2).

The expansion of PK1 and the resultant extended stem is due to a large insertion into the stretch of nucleotides that make up the ring structure (Fig. 3E). In the class 1 xrRNA, the length of the ring is conserved at ∼15 nucleotides, forming a continuous and tight loop around the 5′ end of the xrRNA^5,6,8^. In contrast, in the POWV class 2 the large insertion between P2 and P3 creates an apparent break in the ring at the point of the insertion (Fig. 3D, E). However, closer examination reveals that the ring is still formed, but is now expanded such that it wraps around the 5′ end twice (Fig. 5A). Thus, whereas all other xrRNA structures obtained to date show only a single loop around the 5′ end, the POWV class 2 forms a remarkable ‘double loop’ ring with a full helical turn of base-paired RNA embedded within it.

## Discussion

xrRNAs are ubiquitous in several genera of the *Flaviviridae* family, and are responsible for production of non-coding viral sfRNAs. Understanding of the RNA structure-based mechanism of exoribonuclease resistance has come from solved structures of flavivirus class 1 xrRNAs, but this understanding lacked information on the 3-D structure of the class 2 xrRNAs. Here, we demonstrate the production of sfRNAs in POWV due to the presence of authentic xrRNAs, verified their secondary structures, and solved the 3-D structure of a POWV xrRNA. The structure reveals critical and unique features of the class 2 xrRNAs and completes our understanding of the structural diversity of flavivirus xrRNAs.

One of the most distinctive features of the class 2 xrRNAs compared to previously solved xrRNAs is the extended PK1 and the associated expanded ‘double loop’ ring structure (Fig. 5A). In other solved flavivirus xrRNA structures, the ring wraps around the 5′ end of the RNA only once, using a loop ∼15 nucleotides in length. This is also true in xrRNAs from some plant-infecting viruses^18,19,21^. The purpose of this expansion in the class 2 is not clear, however it is possible that the folding pathway is influenced by the extended PK1; that is, it helps prevent misfolding. However, xrRNAs with a smaller PK1 and ring structure can fold correctly. One other possibility is that the extended structure has a function beyond exoribonuclease resistance, such as binding a specific protein needed for other functions of the 3′UTR. Finally, there may be some advantage to a ‘double loop’ ring around the 5′ end of the xrRNA that may change the mechanism or tune the efficiency of exonuclease resistance.

The presence of the extended ‘double loop’ ring structure in the class 2 xrRNAs raises questions of how this affects the mechanism of exoribonuclease resistance compared to other xrRNAs. Unlike the class 1 xrRNAs, where the exoribonuclease halts on an unpaired region of RNA outside of the ring and pseudoknot, in the class 2 the enzyme halts at locations within PK1 in a region that pairs with nucleotides in the ring structure. This location of the enzyme halt site within PK1, knowledge of the structure and behavior of xrRNA, and the POWV xrRNA 3-D structure now allows a mechanistic model to be proposed. Specifically, as the enzyme degrades in a 5′ to 3′ direction, it must unwind part of the PK1 stem. In fact, because ∼5 nts of single stranded RNA are needed to span the distance between Xrn1’s surface and its catalytic center, almost the entire PK1 stem will be unwound when the enzyme reaches its 3′-most halt point. In this state, the full stretch of RNA between P3 and P1, ∼20 nts, will be unpaired but trapped between the enzyme and the xrRNA core, essentially wrapped twice around the 5′ end of the xrRNA. Thus, in the class 2 xrRNAs, the ring itself is remodeled by the enzyme (Fig. 5B).

We predict that the induced steric clashes produced by this remodeling add substantial resistance to the enzyme’s continued movement. Furthermore, the increasing steric clash as the RNA ring is pressed between the enzyme and the xrRNA core could progressively increase resistance, explaining the ‘stuttering effect’ observed when the halt sites were mapped (Fig. 2C,D). This differs from the class 1 in which there is no evidence that parts of the structure are unwound by Xrn1 or that the ring is remodeled, and the halt site is more precise^5,6,8,9^.

The structure of a class 2 xrRNA presented here, combined with previous subclass 1a and subclass 1b structures, provides a picture of xrRNA diversity in the known flavivirus xrRNAs. All use a similar architecture comprising a central helical junction and two interwoven pseudoknots that create a compact fold. The ring-like structure is also ubiquitous, although the nature of the ring differs between the class 1 and 2, suggesting that different interactions between the ring and the exoribonuclease provide resistance.

Despite the overall similarities in the architecture and the core fold, there are conserved differences that show how each class has evolved a precise and specific set of intramolecular tertiary contacts that cannot be interchanged easily. For example, while all form a ring-like element, the interactions between the base of the P1 stem and L3 that close the ring are different in each class. Likewise, the identity and presence of a nucleotide between P2 and P3, and the overall compactness of the ring element, are different across the classes and are not readily interchanged. Hence, evolution has crafted a set of related RNA folds using different sets of interactions to create similar 3-D outcomes, demonstrating the remarkable structural plasticity and adaptability of RNA structure.

## MATERIALS AND METHODS

### Phylogenetic tree construction

The phylogenetic tree for the described Flaviviridae viruses were constructed using the published maximum likelihood data using the viral NS5, the viral RdRp, described in Bamford et al.^43^ and Mifsud et al.^44^ The tree was rendered in Figtree v1.4.4 (https://github.com/rambaut/figtree/releases) and adjusted in Adobe Illustrator.

### Cell Culture and POWV growth

Vero (African green monkey kidney fibroblast; ATCC CCL-81; passage 15) were grown in 1x MEM (Corning) supplemented with 1% MEM non-essential amino acids (Gibco), 1% Sodium Pyruvate (Gibco), 1x Pen/Strep (Gibco), 1mM HEPES (Gibco), 10% heat inactivated FBS. BHK-21 (baby hamster kidney fibroblast cells; ATCC ccl-10) were maintained in DMEM (Cytiva) supplemented with 1mM sodium pyruvate (Gibco), 27mM sodium bicarbonate (Gibco), 1x Pen/strep (Gibco), and 1uM amphotericin B (Gibco), and 8% heat inactivated FBS.

### POWV stock generation and quantification

All the following steps were performed in a BSL-3 facility. One aliquot of POWV of unknown titer was blindly passaged onto an 80% confluent T-175 flask of Vero cells and incubated for 10 days at 37°C with intermittent rocking. Cellular debris were first clarified from virus containing supernatant by centrifugation at 300 x g for 20 minutes at 4°C. From the clarified supernatant, concentrated viral stocks were made via ultracentrifugation. Briefly, 20 mL of supernatant was gently layered on top of 10mL of 20% sucrose dissolved in PBS (4°C, sterile filtered) in an ultra-clear 1×3.5′ centrifuge tube (Beckman 344058). The tubes were spun down in a SW28 rotor (Beckman) at 27,000rpm for 3.5 hours at 4°C. The supernatant was aspirated, and the pellet was resuspended in complete DMEM + 10% heat inactivated FBS. Concentrated viral stocks were quantified by a plaque-forming unit assay with BHK-21 cells.

### POWV infection in cell culture and total RNA extraction

Vero cells in a 6-well culture plater were inoculated with 0.5mL of POWV stock or mock inoculum at a MOI of 1 or 5 and incubated at 37°C for 1 hour. The inoculum was then aspirated from the cells, washed with 1x PBS, and overlayed with complete 1x MEM. The cells were incubated at 37°C for 24 or 48 hours. After incubation approximately 1.0 x 10^6^ cells per condition were washed 3x in in cold PBS, scraped in buffer TLK (Omega Bio-tek), and homogenized in the Qiashredder (Qiagen) spin column disrupter. Total RNA was purified from the cellular lysate with E.Z.N.A Total RNA Kit (Omega Bio-tek) following the manufacturer’s instructions.

### Northern Blot analysis of POWV sfRNA

For each condition, POWV and mock infection, 1.0 µg of total RNA was resuspended in 2x formamide loading buffer (Thermo) and run on a precast 6% PAGE gel (Invitrogen) with the Riboruler RNA ladder (Thermo). The gels were stained with ethidium bromide and imaged on a UV imager. The RNA was electroblotted to a sheet of HyBond-N^+^ nylon membrane (GE) using the Genie Electrophoretic Transfer Apparatus (Idea Scientific) at 1.6V for 2 hours following the manufactures instructions. UV crosslinking of the RNA to the nylon membrane was performed with a stratalinker (Stratagene) using the “optimal crosslink” setting. Next, the membrane was cut separating the regions containing the sfRNA and the U6 RNA, which were then transferred into individual glass vials and blocked using the ULTRAhyb Oligo Hybridization Buffer (Ambion) at 42°C rotating for two hours.

During the blocking step, the radiolabeled DNA probes were prepared. 100 pmol of POWV_Probe and U6_probe DNA probes were radioactively labeled with 5mCi of [γ^32^-P] ATP using T4 polynucleotide kinase (NEB) for 2 hours at 37°C, and the radiolabeled probes were purified with P-30 spin columns (BioRad). The probes were then heated to 90°C for 2 minutes and quickly resuspended in 10mL of 42°C ULTRAhyb Oligo Hybridization Buffer (Ambion).

The blocking ULTRAhyb Oligo Hybridization Buffer (Ambion) was removed from the membranes and replaced with the 10 mL buffer containing the radiolabeled probe and incubated overnight with rotation at 42°C. The blots were then washed 4 times in 2x saline-sodium citrate (SSC buffer; 300mM NaCl, 30mM Na_3_C_6_H_5_O_7_, 0.5% SDS, pH 7.0) for 15 minutes each at 42°C. The blots were imaged on the Typhoon 9400 scanner (GE Life Sciences) and aligned with the ethidium stained UV image to get the appropriate sizing of the bands.

### Class 2 xrRNA bioinformatic searches and covariance model analysis

An initial class 2 xrRNA alignment was created from a total of 28 sequences of proposed class 2 xrRNA from TBFV 3′ UTR. The sequences were manually aligned with *RALEE* v.0.8^45^ using previous TBFV class 2 secondary structure alignments as a reference. Using *Infernal* v.1.1.4^41^ a database of all available +ssRNA viral sequences downloaded from the National Center for Biotechnology Information (NCBI; last retrieved on 1/06/2023) was searched using default parameters. Hits from the Infernal searches were manually added to the comparative sequence alignment, if they met all the following criteria: *Infernal* E value <0.05, >15% nucleotide variation within each sequence, and location within the 3′ UTR. Sequences with higher E values were also inspected and added to the list if they met the last three requirements. For the final proposed covariance model of 36 class 2 xrRNA sequences, statistical validation was performed using *R-scape* v2.0.0.j^37,40^ and rendering the covariance model with *R2R* v.1.0.5^42^.

### *In vitro* RNA transcription

DNA templates were ordered as gBlock DNA fragments (IDT) and cloned into pUC19. 200 µL PCR reactions using primers containing an upstream T7 promoter were used to generate dsDNA templates for transcription. Typical PCR conditions: 100 ng plasmid DNA, 0.5 µM forward and reverse DNA primers (Supplementary Table 2), 500 µM dNTPs, 25 mM TAPS-HCl (pH 9.3), 50 mM KCl, 2 mM MgCl_2_, 1 mM β-mercaptoethanol, and Phusion DNA polymerase (New England BioLabs). Templates for RNA used in aminoacylation assays were amplified using reverse primers containing two 5′-terminal 2′-*O*-methyl modified bases to ensure the correct 3′ end of the RNA. dsDNA amplification was confirmed by 1.5% agarose gel electrophoresis. Transcriptions were performed in 1 mL volume using 200 µL of PCR product (∼ 0.1 µM template DNA) and 10 mM NTPs, 75 mM MgCl_2_, 30 mM Tris-HCl (pH 8.0), 10 mM DTT, 0.1% spermidine, 0.1% Triton X-100, and T7 RNA polymerase. Reactions were incubated at 37°C overnight. After transcription, insoluble inorganic pyrophosphate was removed by centrifugation at 5000xg for 5 minutes, then the RNA-containing supernatant was ethanol precipitated with 3 volumes of 100% ethanol at −80°C for a minimum of 1 hour and then centrifuged at 21000xg for 30 minutes at 4°C to pellet the RNA, and the ethanolic fraction was decanted. The RNA was resuspended in 9 M urea loading buffer then purified by denaturing 10% PAGE. Bands were visualized by UV shadowing then excised. Bands were then crush-soaked in diethylpyrocarbonate-treated (DEPC) milli-Q water at 4°C overnight. The RNA-containing supernatant was then concentrated using spin concentrators (Amicon) to the appropriate concentration in DEPC-treated water. RNAs were stored at −80°C with working stocks stored at −20°C.

### Expression of Kluyveromyces lactis Xrn1ΔC

The DNA sequence encoding the Xrn1 enzyme from *K. lactis* (residues 1-1245) containing an in-frame C-terminal hexahistidine affinity tag was a kind gift from Dr. Liang Tong at Columbia University. The protein was recombinantly expressed in BL21 (DE3) cells. Cells were grown in LB to an OD_600_ of 0.6, then protein expression was induced using 500 µM isopropyl β-D-1-thiogalactopyranoside (IPTG) overnight at 18°C. Pelleted cells were resuspended in lysis buffer containing 20 mM Tris-HCl (pH 7.0), 500 mM NaCl, 2 mM β-mercaptoethanol (BME), 10% (v/v) glycerol, and cOmplete EDTA-free protease inhibitor cocktail tablets (Roche). Cell lysate was then sonicated on ice for 2 minutes of: 20 seconds on, 40 seconds off at 75 W. Cell lysate was clarified by centrifugation at 30,000xg for 30 minutes at 4°C. The soluble fraction was purified by nickel affinity chromatography in a buffer containing 150 mM NaCl, 20 mM Tris-HCl (pH 7.0), 200 mM imidazole, 10% glycerol, and 2 mM β-mercaptoethanol. The protein was exchanged into a storage buffer containing 50 mM Tris-HCl (pH 8.0), 150 mM NaCl, 2 mM DTT, and 50% glycerol using a spin concentrator (Amicon) and stored at 1.0 mg mL^-1^ at −80°C with working stocks stored at −20°C.

### Expression of Bdellovibrio bacteriovorus RppH

A plasmid containing BdRppH (Uniprot ID Q6MPX4_BDEBA, Q6MPX4) in frame with a hexahistidine tag was a kind gift of J. Belasco at New York University. BdRppH was purified in a manner identical to that of Xrn1ΔC as described above. The protein was concentrated to 12 mg mL^−1^ in a buffer containing 150 mM NaCl, 25 mM Tris (pH 7.5), 1 mM DTT, and 50% (v/v) glycerol and then stored at −80 °C.

### *In vitro* RNA degradation assays

Assays were performed using 2 µg of RNA in a 20 µL volume. RNA was refolded as described above the divided into two 10 µL volumes. To both 1 µL of BdRppH was added and to one 1 µL of KlXrn1ΔC was added. To the other volume 1 µL of KlXrn1ΔC storage buffer (150 mM NaCl, 50 mM Tris, 2 mM DTT, 50% (v/v) glycerol, pH 8.0) was added. The reactions were then incubated at 37°C for 2 hours. The reactions were quenched with the addition of 1 volume of 9 M urea loading buffer and resolved on a 10% denaturing PAGE gel then the RNA was visualized by methylene blue staining

### Stop site analysis

Full length POWV 3′UTR was degraded with Xrn1 *in vitro* following the *in vitro* Xrn1 degradation assay outlined above. First, a FAM labeled primer specific to the 3′ terminal end was annealed to the RNA and then immobilized on oligo-dT_25_ beads (Invitrogen). The RNA was reverse transcribed on the oligo-dT_25_ beads with 2.5 µL of reverse transcription mixture of 1.0 µL of 5x First strand Buffer (Invitrogen), 0.25 µL of 0.1 M 1,4-Dithiothreitol (DTT), 0.4 µL of a 10 mM equimolar mixture of dNTPs, 0.75 µL of DEPC-Treated H_2_O, and 0.1 µL of Superscript III reverse transcriptase (Invitrogen). The mixture was incubated at 52°C for 50 min and the RNA was subsequently hydrolyzed with 5 µL of 0.4 M NaOH at 65°C for 20 min. The samples were cooled on ice for 2 min and then neutralized with 5 µL of an acid quench mixture consisting of 1.4 M NaCl, 0.6 N HCl, and 1.3 M NaOAc. The supernatant was aspirated from each well, and the bound complementary DNA (cDNA) was washed three times with 100 µL of 70% EtOH. The beads were then dried for 15 minutes to ensure that there was no residual EtOH present. The cDNA was eluted in 11 µL of a ROX-formamide mixture composed of 2.75 µL of ROX-500 ladder (Applied Biosystems) in 1.2 mL of HiDi-formamide (Applied Biosystems) at room temperature for 15 min. The 11 µL cDNA mixture was transferred to an optical plate and analyzed with an ABI 3500 Genetic Analyzer Capillary Electrophoresis machine (GE).

In parallel, a Sanger sequencing ladder was constructed through reverse transcription of unmodified POWV 3′UTR (Table S2) in the presence of dideoxynucleoside triphosphates (ddNTP′s). In brief 2.5 µL of an RNA mixture was made consisting of 0.2 µL of 0.25 µM FAM primer, 1.5 µL of oligio-dT_25_ beads (Invitrogen), 0.5 µL of 2.4 µM RNA, and 0.25 µL of 5 M betaine. To this, 2.5 µL of a ladder reverse transcription mixture were added consisting of 1.0 µL of 5x First strand Buffer (Invitrogen), 0.25 µL of 0.1M DTT, 0.4 µL equimolar mix of 1.0 mM dNTP′s, 0.4 µL of 1.0 mM of respective ddNTP, 0.1 µL of Superscript III reverse transcriptase (Invitrogen), and 0.25 µL of 5 M Betaine. The cDNA was eluted off the beads in 11 µL of a ROX-formamide outlined above. The 11 µL cDNA mixture was transferred to an optical plate and analyzed with an ABI 3500 Genetic Analyzer Capillary Electrophoresis machine (GE). After running the xrRNA degraded 3′UTR and the Sanger Sequencing ladder on the ABI 3500 Genetic analyzer Capillary Electrophoresis machine, the HiTRACE MATLAB suite^46^ (https://ribokit.github.io/HiTRACE/) with MATLAB (v8.3.0.532) was used to analyze the stop site data.

### *In vitro* probing of POWV 3’UTR, sfRNA1, and sfRNA2

#### 1m7 and DMS probing of the POWV 3’UTR

1.0 µg of *in vitro* transcribed full length POWV 3′UTR was heat denatured in 12 µL DEPC treated H_2_O at 95°C for two minutes and then immediately cooled on ice. To this 8.0 µL of concentrated RNA folding buffer was added to a final concentration of 100 mM NaCl, 50 mM Tris pH 7.5, 4 mM MgCl_2_, and folded for 20 minutes at 55°C. The folded RNA was slow cooled to room temperature. After cooling the reaction was split into two, and 0.5 µg of the RNA was labeled with 10mM 1m7 (Sigma) or DMSO (Sigma) at 37°C for 75 seconds (five 1m7 hydrolysis half-lives). In parallel, 0.5 µg of folded RNA was modified with 0.5% DMS (Sigma) or an equal amount of EtOH as a negative control for 5 minutes at 37°C, and then quenched with 2-Mercaptoethanol (Sigma). All conditions were purified and concentrated using the RNeasy Micro Kit (Qiagen).

#### 1m7 and DMS probing of the POWV sfRNA1 and sfRNA2

2 µg of folded *in vitro* transcribed POWV 3′UTR was treated with Xrn1 following the methods outlined above. After the 2 hour incubation, DMS, 1m7, or their appropriate controls was added to the reaction at the same final concentrations as the 3’UTR probing outlined above. The reactions were cleaned up using the RNeasy Micro Kit (Qiagen) and separated on a 6% denaturing PAGE gel. The sfRNA1 and sfRNA2 bands were visualized by UV shadowing, cut from the gel, and eluted at 4°C overnight in ∼ 5 mL 50 mM Sodium cacodylate pH 6.5, 0.1 mM EDTA, and 50mM NaCl. The RNA was concentrated with a 30K molecular weight cut off Amicon Ultra spin Concentrators (Millipore-Sigma) and buffer exchanged into DEPC water.

#### Reverse transcription and second strand synthesis

The following protocol was applied to the full length 3′UTR, sfRNA1 and sfRNA2 RNA species. Approximately 400 ng of modified or control RNA was incubated with of 200 ng/µL random nonamer primer (NEB; S1254S) and 1.0 µL of POWV_SHAPEMAP_RT_Primer (Supplemental Table S2) at 65°C for 5 minutes and then cooled on ice. To each of the reactions 8.0 µL of MAP buffer (125mM Tris pH 8.0, 187.5mM KCl, 15mM MnCl_2_, 25mM DTT, 1.25mM equimolar dNTP mix, and 200U Superscript III) was added to the reaction and incubated at 25°C for 2 minutes. The reaction was then incubated at 25°C for 10 minutes, 42°C for 3 hours, 70°C for 15 minutes, and cooled to 4°C. The RNA/cDNA hybrids were then purified on the G25 (Cyteva) spin column following the manufacturers protocol. Second strand synthesis of the cDNA was carried out using the NEBNext second strand synthesis kit (NEB) following the manufacturer’s instructions. The resultant cDNA was purified using the DNA Clean & Concentrator-5 kit (Zymo) eluting in 10 µL of DEPC-treated H_2_O and quantified fluorometrically with the Qubit (Thermo) using the dsDNA HS Assay Kit (Thermo).

#### Library prep and sequencing

Next generation sequencing library prep was performed with the Nextera XT DNA library preparation kit (Illumina) using the Nextera XT Index Kit v2 Set B (Illumina). Samples were normalized and pooled together using a combination of the Qubit (Thermo) and the 4200 TapeStation System (Agilent Technologies) with the High Sensitivity D1000 Screen Tape (Agilent Technologies). Final libraries were sequenced on the NovaSeq 6000 platform (Illumina).

#### Data analysis

Raw sequencing reads were first assessed with FastQC and adapter trimmed using Trimmomatic v0.40^47^. The processed reads were then analyzed with Shapemapper2^48^ with the default parameters and a minimum read depth of 5000 reads. The resultant data was analyzed using a custom in house pipeline of Python v3.9 and R v4.2.2 scripts. Reactivities were then plotted onto the proposed secondary structure using Varna 3.93^49^ and plots were constructed using GGplot2 v3.4.0 and R v4.2.2.

### In Vitro RNA transcription and purification for Cryo-Electron Microscopy

DNA templates were ordered as gBlock DNA fragments (IDT) and cloned into pUC19 or as overlapping primers. 1 mL PCR reactions using primers containing an upstream T7 promoter were used to generate dsDNA templates for transcription. Typical PCR conditions: 1 ng/µL plasmid DNA, 0.5 µM forward and reverse DNA primers, 500 µM dNTPs, 25 mM TAPS-HCl (pH 9.3), 50 mM KCl, 2 mM MgCl_2_, 1 mM β-mercaptoethanol, and Phusion DNA polymerase (New England BioLabs). dsDNA amplification was confirmed by 1.5% agarose gel electrophoresis. Transcriptions were performed in 5 mL volume using 1 mL of PCR product (∼ 0.1 µM template DNA) and 10 mM NTPs, 75 mM MgCl_2_, 30 mM Tris-HCl (pH 8.0), 10 mM DTT, 0.1% spermidine, 0.1% Triton X-100, and T7 RNA polymerase. Reactions were incubated at 37°C for 3 hours. After transcription, insoluble inorganic pyrophosphate was removed by centrifugation at 5000xg for 5 minutes, then the RNA-containing supernatant was ethanol precipitated with 3 volumes of 100% ethanol at −80°C for a minimum of 1 hour and then centrifuged at 21000xg for 30 minutes at 4°C to pellet the RNA, and the ethanolic fraction was decanted. The RNA was resuspended in 9 M urea loading buffer then purified by denaturing 6% PAGE. Bands were visualized by UV shadowing then excised. Bands were then crush-soaked in diethylpyrocarbonate-treated (DEPC) milli-Q water at 4°C overnight. The RNA-containing supernatant was then concentrated using spin concentrators (Amicon) to the appropriate concentration in DEPC-treated water. RNAs were stored at −80°C with working stocks stored at −20°C.

### POWV RNA refolding

RNA was refolded at the described concentrations as follows. To the RNA solution (usually 18 µL of 3 mg/mL), 1/20^th^ the volume of 1 M Tris pH 7.5 was added. The solution was heated to 90°C for 3 minutes then allowed to cool to room temperature for 10 minutes. Subsequently, 1/20^th^ the volume of 200 mM MgCl_2_ was added. The solution was then heated to 50°C for 30 minutes then allowed to cool to room temperature for 10 minutes. Once cooled, the folded RNA solution was stored on ice until use.

### Cryo-EM sample preparation and data acquisition

RNA sample was applied to a glow discharged C-flat (1.2/1.3, 400 mesh) holey carbon grid. Samples were vitrified in liquid ethane using a Vitrobot Mark IV (5 s wait time, 5.5 s blot, −5 blot force, 100% humidity, 4°C). Samples were imaged on a Titan Krios equipped with a Gatan K3 camera and Bioquantum energy filter. Movies were collected in fringe free mode, with a physical pixel size of 0.788 Å, total dose of 50 e^-^/Å^2^, and a defocus range of −0.8 to −2.0 µm. SerialEM^50^ was used for all data collection. Additional information on data collection can be found in Supplementary Table 1.

### Cryo-EM data processing

The general data processing workflow was approximately equivalent for all datasets using cryoSPARC 4.0. Briefly, all data was patch motion corrected and the contrast transfer function was locally fit. An initial set of particles was picked using blob picker from the first 1000 micrographs. Particles were extracted and 2D classified. Good classes for template picking were manually chosen. Template based picking was performed using template picker on the entire dataset. Particles were then extracted and subjected to several rounds of 2D classification to sort out junk particles. Good 2D classes and the associated particles were manually curated after each round. The pruned particles were then used for initial map generation using the Ab Initio reconstruction job requesting 2 volumes. The resulting 2 volumes were used for heterogenous refinement, where one class was the “good” class and the other a “junk” class to remove poor quality particles. Several rounds of heterogenous refinement were performed, discarding the “junk” particles after each round until the map quality stopped improving. Subsequently, the “good” volume and particles were subjected to non-uniform refinement. Local motion correction, local CTF refinement, and global CTF refinement were then carried out on the data and the particles were subjected to further rounds of non-uniform refinement to yield the final maps. Additional information on data processing for each map can be found in Supplementary Figure 4.

### Structure modeling

To build the atomic model, the Cryo-EM structure of the tetrahymena ribozyme was placed into the maps using ridged body fitting (PDB: 8TJX)^31^ followed by the solved crystal structure of the Tamana bat virus xrRNA (PDB:7K16)^8^. The model was then manually built using the TABV xrRNA as an initial reference and joined in COOT^51^. The final model was refined against the map by way of several rounds of real-space refinement in PHENIX^52^ and manual adjustments in COOT. The statistics of model refinement and validation are listed in Supplementary Table 1.

## ACKNOWLEDGEMENTS

The authors thank current and past members of the Kieft Lab for critical reading of the manuscript and insightful comments. Theo Humphries (PNCC) assisted with microscope operations. David Beckham (UT Southwestern) provided helpful discussions for the BSL3 work performed herein. This work was supported by R01AI133348 to J.S.K. A portion of this research was supported by NIH grant U24GM129547 and performed at the PNCC at OHSU and accessed through EMSL (grid.436923.9), a DOE Office of Science User Facility sponsored by the Office of Biological and Environmental Research.

