## Supplemental Figures and Tables for "Tick-borne flavivirus exoribonuclease-resistant RNAs contain a ‘double loop’ structure"

### Tick-Borne Flaviviruses Uses a Structurally Distinct RNA to Prevent Exonuclease Degradation

**Supplementary Figure 1:** 1m7 probing of the POWV 3'UTR, sfRNA1, and sfRNA2
**Supplementary Figure 2:** DMS probing of the POWV 3'UTR, sfRNA1, and sfRNA2
**Supplementary Figure 3:** Pseudoknot mutations tested
**Supplementary Figure 4:** POWV xrRNA1 cryo-EM construct and validation
**Supplementary Figure 5:** Scaffolded POWV xrRNA1 cryo-EM processing workflow
**Supplementary Figure 6:** Class 2 xrRNA comparative sequence alignment
**Supplementary Figure 7:** Alternative depictions of class 2 xrRNA consensus secondary structure
**Supplementary Table 1:** Cryo-EM data collection, refinement, and validation statistics
**Supplementary Table 2:** Oligos used in this study

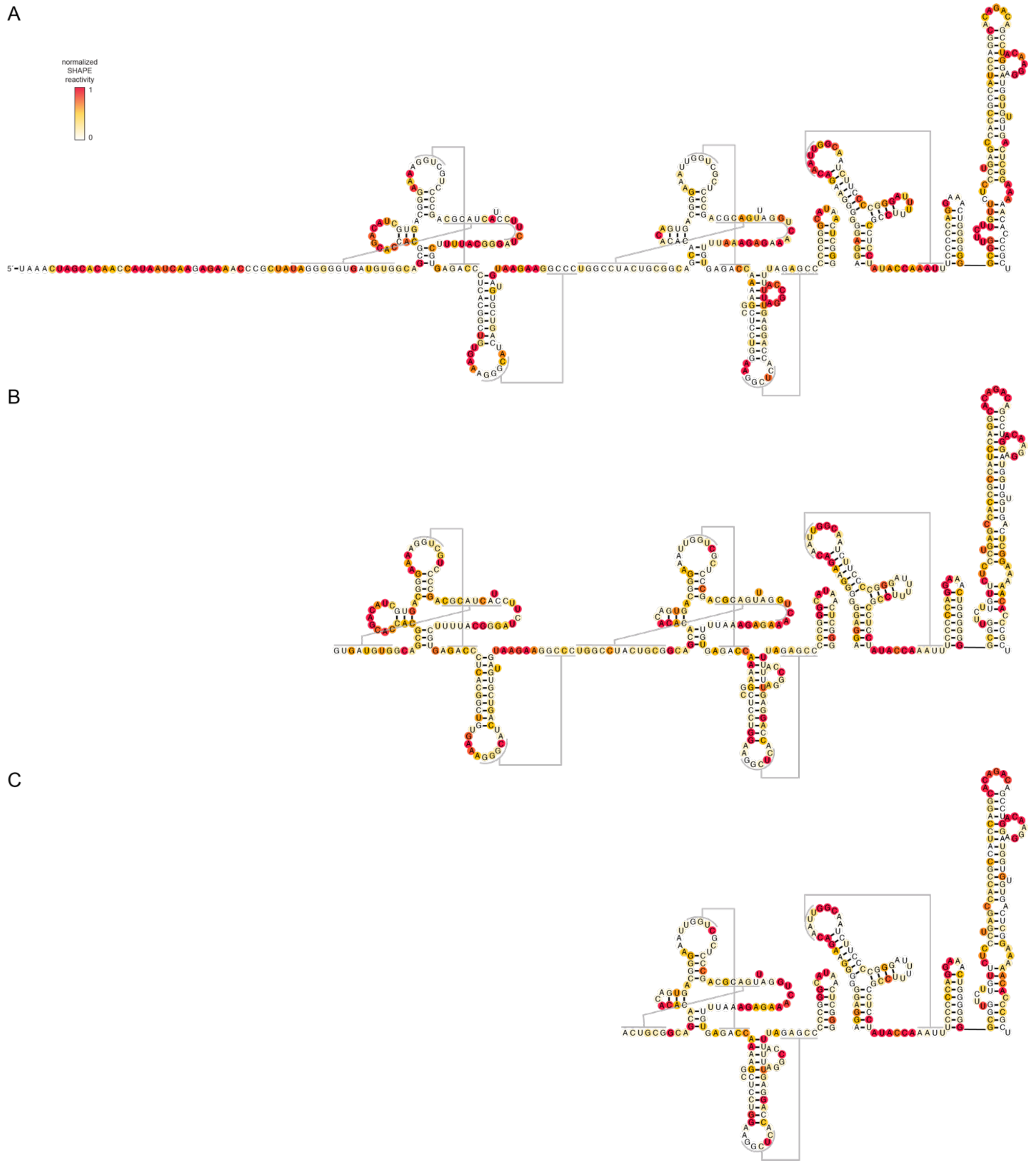


**Supplementary Figure 1. 1m7 probing of the POWV 3'UTR, sfRNA1, and sfRNA2.**
1m7 chemical probing of the POWV 3'UTR (**A**), sfRNA1 (**B**), and sfRNA2 (**C**). In the case of B and C, *in vitro* transcribed RNAs were treated with XRN1 *in vitro* then purified and chemically probed using 1m7. Reactivities are averages of a minimum of three experiments. Reactivities are mapped onto the bioinformatically supported secondary structure of the 3'UTR.

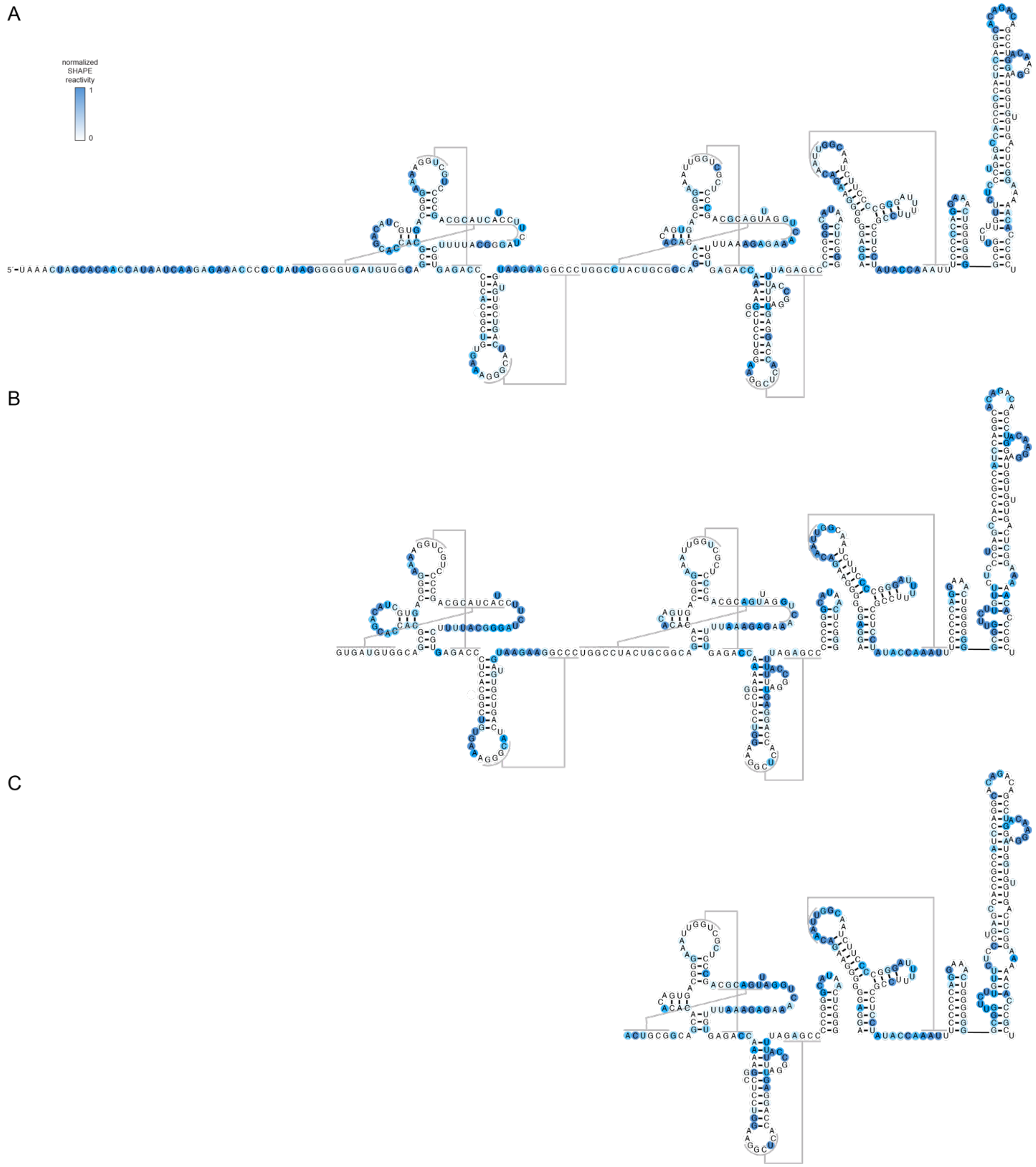


**Supplementary Figure 2. DMS probing of the POWV 3'UTR, sfRNA1, and sfRNA2.**
DMS chemical probing of the POWV 3'UTR (**A**), sfRNA1 (**B**), and sfRNA2 (**C**). In the case of B and C, *in vitro* transcribed RNAs were treated with XRN1 *in vitro* then purified and chemically probed using DMS. Reactivities are averages of a minimum of three experiments. Reactivities are mapped onto the bioinformatically supported secondary structure of the 3'UTR.

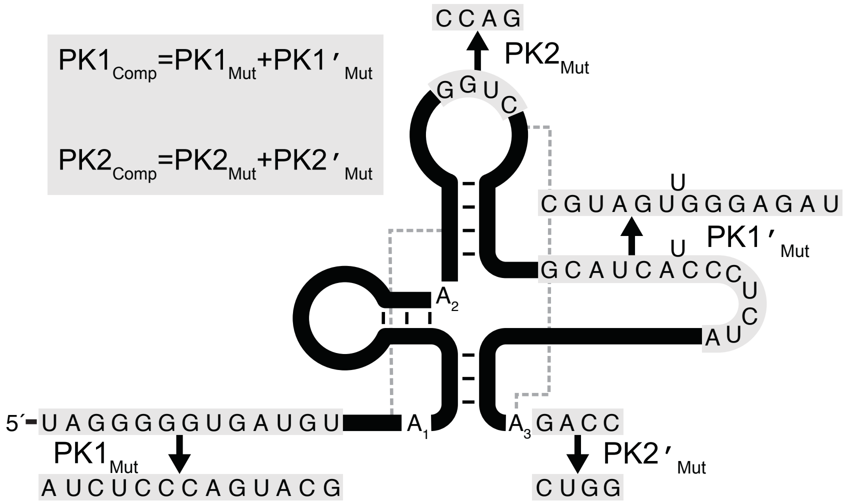


**Supplementary Figure 3. Pseudoknot mutations tested.**
Graphical description of pseudoknot mutations tested in the context of the POWV xrRNA1. Single mutations were made as shown and double mutations of each pseudoknot were made with the combinations described in the inset.

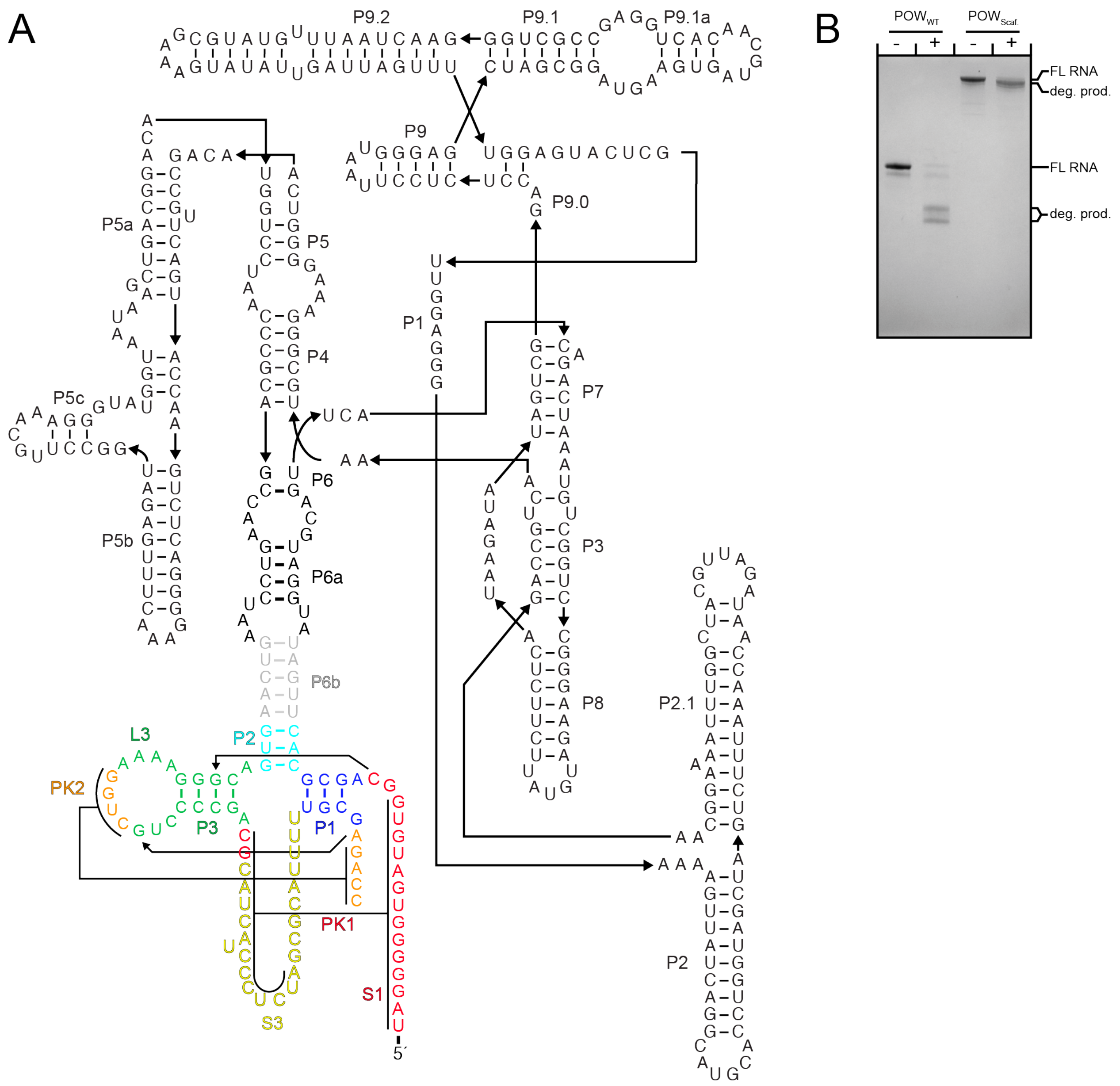


**Supplementary Figure 4. POWV xrRNA1 cryo-EM construct and validation.**
**A.** Secondary structure of the POWV xrRNA1 RNA grafted onto a circularly permuted tetrahymena ribozyme for cryo-EM structure determination. The POWV xrRNA1 portion is colored corresponding to figure 3. **B.** In vitro Xrn1 degradation assay of the WT POWV xrRNA1 RNA and the scaffolded POWV xrRNA1 RNA demonstrating the construct retains resistance and as such is correctly folded.

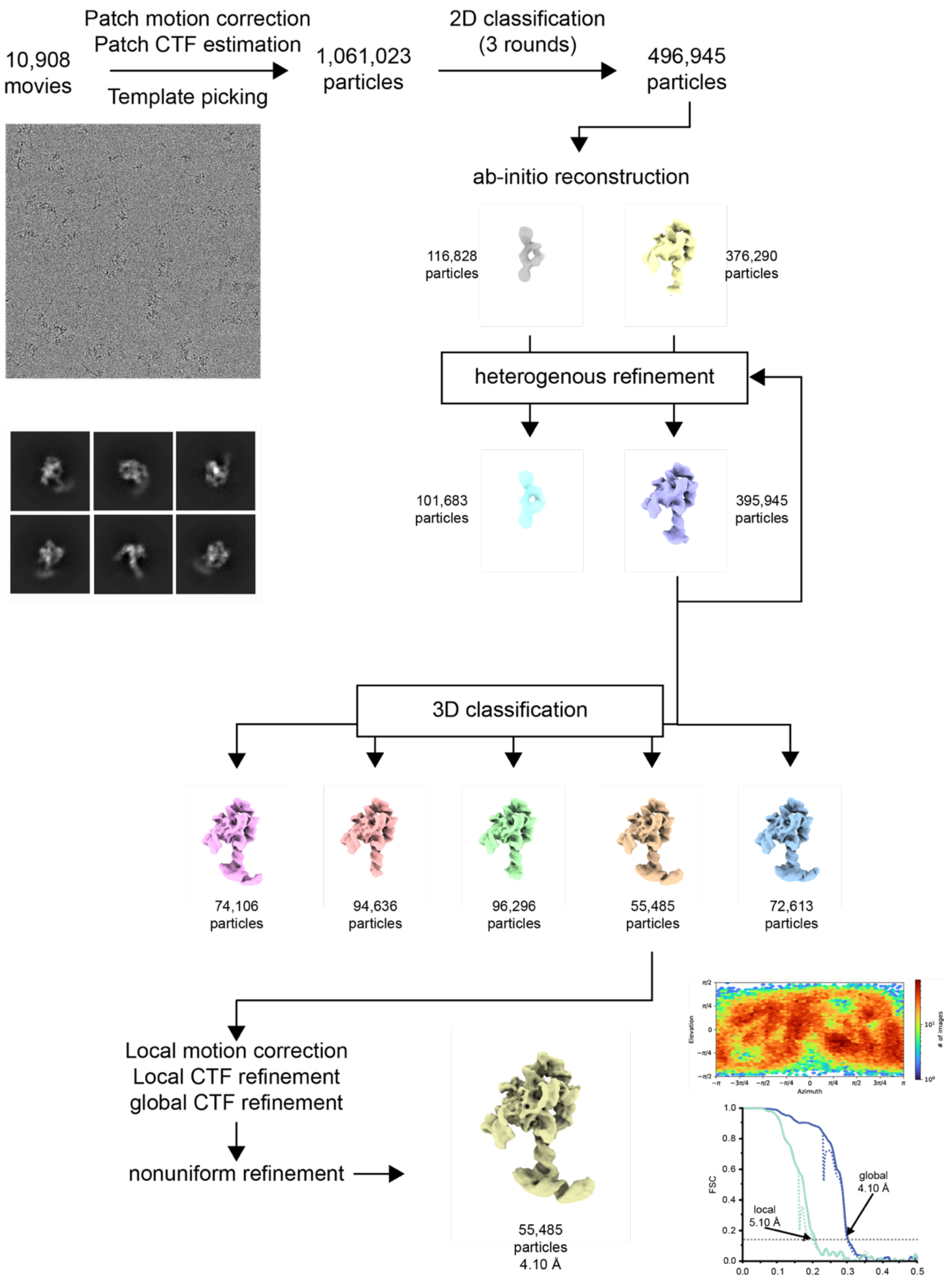


**Supplementary Figure 5. Scaffolded POWV xrRNA1 cryo-EM processing workflow.**
Cryo-EM processing workflow of the scaffolded POWV xrRNA1 in cryoSPARC. A representative micrograph and 2D classes are shown. The Viewing orientation distribution and FSC graph for global and xrRNA local resolution are shown for the final model used for model building.

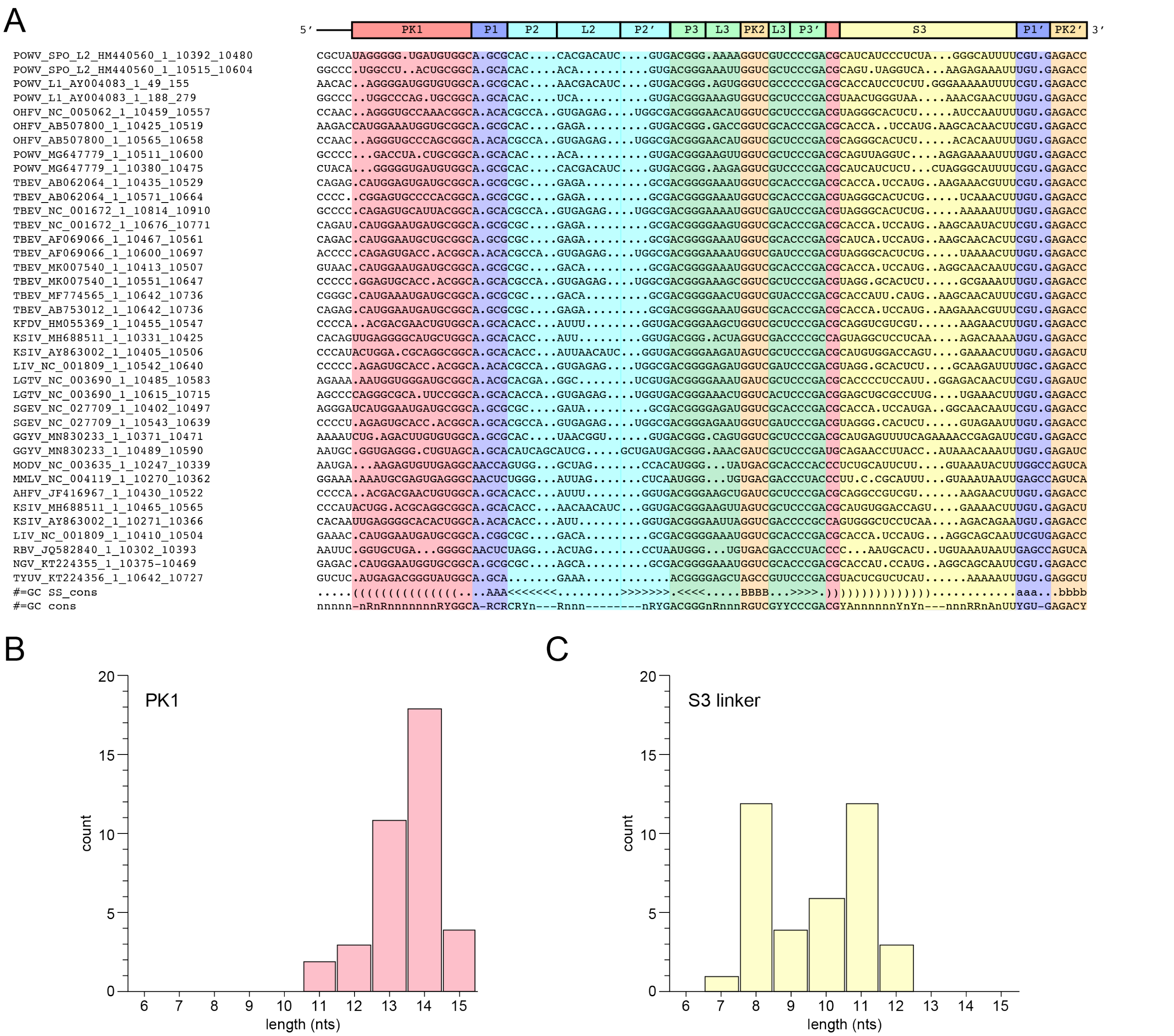


**Supplementary Figure 6. Class 2 xrRNA comparative sequence alignment.**
**A.** Alignment of the 38 class 2 xrRNAs identified in our bioinformatic searches aligned based on conserver secondary structure. Regions are colored corresponding to figure 3 for simplicity. **B.** Length distribution of PK1 in alignment. **C.** Length distribution of S3 linker (unpaired region) in alignment.

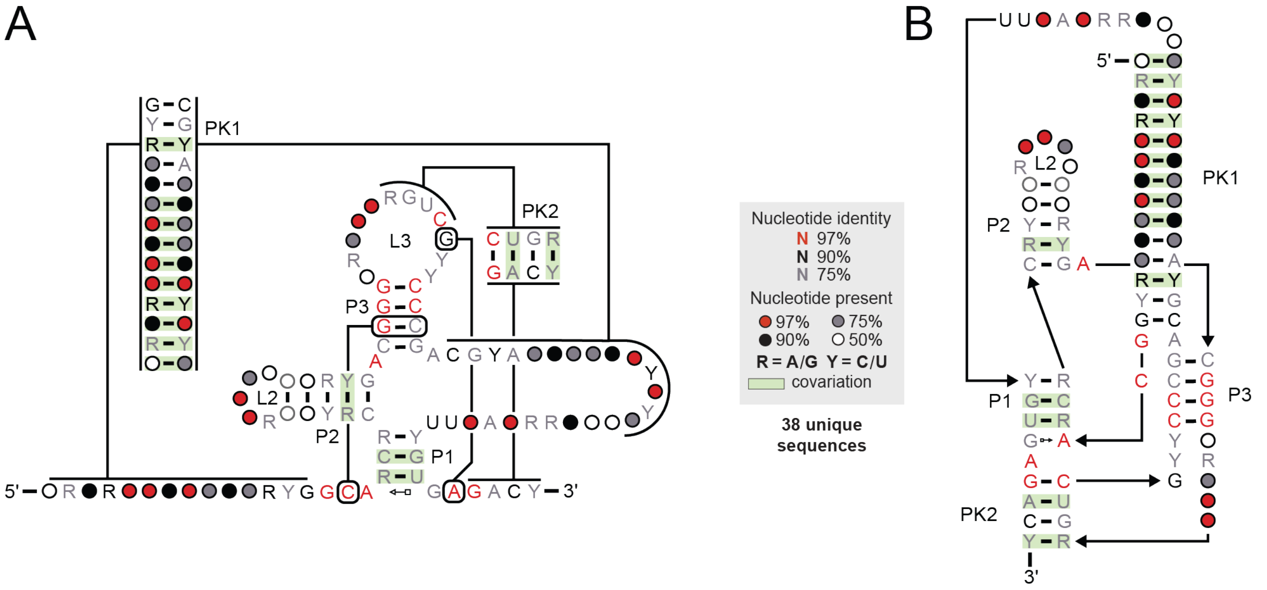


**Supplementary Figure 7. Alternative depictions of class 2 xrRNA consensus secondary structure.**

**A.** Consensus secondary structure of class 2 xrRNA depicted as drawn in figure 3C and 4A. **B.** Consensus secondary structure of class 2 xrRNA drawn in an alternative depiction, though maintaining the same contact, to emphasize the critical tertiary contact present in this class.

| | TetP6b + POWV xrRNA (EMD-xxx) (PDB xxx) |
|---|---|
| **Data collection and processing** | |
| Grid Type | C-flat 1.2/1.3 |
| Microscope | FEI Titan Krios |
| Camera | K3 |
| Magnification | 105,000 |
| Voltage (kV) | 300 |
| Electron exposure (e–/Å$^2$) | 50 |
| Defocus range (μm) | -0.8 to -2.0 |
| Pixel size (Å) | 0.788 |
| Symmetry imposed | C1 |
| Initial particle images (no.) | 1,061,023 |
| Final particle images (no.) | 55,485 |
| Map resolution (Å) | 4.10 |
| FSC threshold | 0.143 |
| Map resolution range (Å) | 4.010-8.233 |
| **Refinement** | |
| Initial model used (PDB code) | 8TJX |
| Model resolution (Å) | 4.10 |
| FSC threshold | 0.143 |
| Mask CC | 0.880 |
| Model resolution range (Å) | 4.0-8.0 |
| Map sharpening *B* factor (Å$^2$) | -135.4 |
| Model composition | |
| Non-hydrogen atoms | 9416 |
| RNA bases | 440 |
| Ligands | 19 |
| *B* factors (Å$^2$) | |
| RNA | 150.22 |
| Ligand | 78.83 |
| R.m.s. deviations | |
| Bond lengths (Å) | 0.003 |
| Bond angles (°) | 0.559 |
| Validation | |
| MolProbity score | 3.43 |
| Clashscore | 4.34 |
| Poor rotamers (%) | 0 |

**Supplementary Table 1. Cryo-EM data collection, refinement, and validation statistics**

**Supplementary Table 2. Oligos used in this study**

| **Oligo** | **Sequence 5' to 3'** |
|---|---|
| POWV 3' UTR gBlock | GGCTATCGAATTCTAATACGACTCACTATAGGACTAGCACAACCATAATCAAGAGAAACCCGCTATAGGGGGTGATGTGG<br>CAGCGCACCACGACATCGTGACGGGAAAAGGTCGTCCCCGACGCATCATCCCTCTAGGGCATTTTCGTGAGACCCTCAC<br>GGCTGTGAAAGGGCATCAGTCGTGTAGTAAGAAGGCCCTGGCCTACTGCGGCAGCACACACAGTGACGGGAAATTGGT<br>CGCTCCCGACGCAGTTAGGTCAAAGAGAAATTTGTGAGACCAAAAGGCCTCCTGGAAGGCTCACCAGGAGTTAGGCCAT<br>TTTAGAGCCCCCGGGCATAACTCGGGAGGAGGGGGGAAGACAATTGGCAATCTTCCCCGGGATTTTTCCGCCTCCTATA<br>CCAAATTTCCCCCAGGAAACTGGGGGGGCGGTTCTTGTTCTCCCTGAGCCACCGCCATCCAGGCACAGACAGCCTGAC<br>AAGGAGATGGTGTGTGACTCGGAAAAACACCCGCTGGATCCTACCGGA |
| POWV 3' UTR | GGACTAGCACAACCATAATCAAGAGAAACCCGCTATAGGGGGTGATGTGGCAGCGCACCACGACATCGTGACGGGAAA<br>AGGTCGTCCCCGACGCATCATCCCTCTAGGGCATTTTCGTGAGACCCTCACGGCTGTGAAAGGGCATCAGTCGTGTAGT<br>AAGAAGGCCCTGGCCTACTGCGGCAGCACACACAGTGACGGGAAATTGGTCGCTCCCGACGCAGTTAGGTCAAAGAGA<br>AATTTGTGAGACCAAAAGGCCTCCTGGAAGGCTCACCAGGAGTTAGGCCATTTTAGAGCCCCCGGGCATAACTCGGGAG<br>GAGGGGGGAAGACAATTGGCAATCTTCCCCGGGATTTTTCCGCCTCCTATACCAAATTTCCCCCAGGAAACTGGGGGGG<br>CGGTTCTTGTTCTCCCTGAGCCACCGCCATCCAGGCACAGACAGCCTGACAAGGAGATGGTGTGTGACTCGGAAAAACA<br>CCCGCT |
| POWV Cryo Construct | GCCGATGAATTCTAATACGACTCACTATAGGACTAGCACAACCATAATCAAGAGAAACCCGCTATAGGGGGTGATGTGGC<br>AGCGCACTTGATATGGATGCAGTTCACAGACTAAATGTCGGTCGGGGAAGATGTATTCTTCTCATAAGATATAGTCGGAC<br>CTCTCCTTAATGGGAGCTAGCGGATGAAGTGATGCAACACTGGAGCCGCTGGGAACTAATTTGTATGCGAAAGTATATTG<br>ATTAGTTTTGGAGTACTCGTTGGAGGGAAAAGTTATCAGGCATGCACCTGGTAGCTAGTCTTTAAACCAATAGATTGCATC<br>GGTTTAAAAGGCAAGACCGTCAAATTGCGGGAAAGGGGTCAACAGCCGTTCAGTACCAAGTCTCAGGGGAAACTTTGAG<br>ATGGCCTTGCAAAGGGTATGGTAATAAGCTGACGGACATGGTCCTAACCACGCAGCCAAGTCCTAAGTCAAGTGACGGG<br>AAAAGGTCGTCCCCGACGCATCGTGAGACCCAGGATCCTACCGGA |
| M13_Forward_PCR_Primer | GTA AAA CGA CGG CCA GTG |
| POWV_Reverse_PCR_Primer | AGCGGGTGTTTTTCCGAGTCACAC |
| POWV_Fam_Labled_Primers | /5–6FAM/AGCGGGTGTTTTTCCGAGTCACAC |
| POWV_Cryo_Construct_Fwrd | TAATACGACTCACTATAGGATAGGGGGTGATGTGGCAGC |
| POWV_Cryo_Construct_Reverse | TGGGTCTCACGATGCGTCGG |
| POWV_SHAPEMAP_RT_Primer | TGGGTCTCACGATGCGTCGG |
| POWV_Probe | GGTATAGGAGGCGGAAAAATCCCGGGGAAGAT |
| U6_Probe | TATGGAACGCTTCACGAATTTGCGTGTCATCC |
